# A new Mmrn1-CreERT2 Mouse Model for High-Fidelity Labeling of Adult Lymphatic Endothelium

**DOI:** 10.64898/2026.10.16.759619

**Authors:** Coby Goelz, Gabriel A Tavares, Joseph Lubera, Antoine Louveau

**Author notes:** equal-contribution (Coby Goelz, Gabriel A Tavares). Lead Contact: Antoine Louveau, PhD, 9500 Euclid Avenue NC30, Cleveland, OH, 44195.

## Abstract

Cell type-specific Cre driver lines are essential for studying organ-specific biology in vivo, but recombination in unintended targets can limit their interpretive power. The dural meninges harbor a distinct network of lymphatic vessels implicated in neurological processes ranging from neurodevelopment to neurodegeneration. Current lymphatic reporters, driven by Prox1, Lyve1 or Vegfr3, efficiently recombine in lymphatic endothelial cells, including in the dura, but are also expressed in other CNS cell populations, resulting in non-lymphatic recombination that complicates interpretation. Here, we identify Mmrn1 as a gene specifically enriched in lymphatic endothelial cells throughout the body, with no detectable expression in other CNS cells. We generated an Mmrn1CreERT2 mouse line and show that, in adult mice, it drives recombination restricted to lymphatic endothelial cells, with no detectable recombination in non-lymphatic CNS populations. A similar but less complete recombination is observed in developing mice. This tool offers the lymphatic field a new means to specifically manipulate lymphatic vasculature and study its cell-specific functions.

## Introduction

The lymphatic vasculature maintains tissue homeostasis by draining interstitial fluid (ISF) and macromolecules, as well as ensuring immune cell transport into lymph nodes^1–4^. Specifically for the central nervous system (CNS), the network of CNS-associated lymphatic vessels has emerged as a central player for brain function in both homeostatic conditions and during diseases^5,6^. Primarily located within the dura mater layer of the meninges, they drain cerebrospinal fluid (CSF), meningeal ISF, and local immune cells into the cervical lymph nodes^7–10^. Dural lymphatic vessels differ from peripheral lymphatics in several aspects: (1) they develop postnatally rather than embryonically, arising first at the skull base and along cranial nerves before extending along the dural sinuses^11–13^ and (2) they remain under the control of the VEGF-C/VEGFR3 pathway signaling past their developmental window^11,12,14^. These differences make the meningeal bed both biologically intriguing and technically demanding to study.

Moving from correlation to causation in this field requires the capacity to delete or report genes selectively in lymphatic endothelial cells (LECs). In practice, the field relies on a small number of Cre drivers built around the canonical LEC markers: Prox1^15,16^, Lyve1^17^, and Flt4/VEGFR3^18–20^. Each of these markers is specific to lymphatics only within a given anatomical context. Prox1 is a transcription factor required for LECs specification^21,22^, but it is also expressed in some hepatocytes^23^, developing cardiomyocytes^24^ and, critically for CNS work, in neuronal populations including dentate gyrus granule cells^25^. Lyve1 is a hyaluronan receptor expressed by LECs^26^ but also by liver and spleen sinusoidal endothelial cells^27^, and by a large population of macrophages, particularly in the CNS^28–30^. Vegfr3 is re-expressed by angiogenic and fenestrated blood endothelium^31–33^. The consequence is that a conditional deletion driven by any of these promoters in the CNS simultaneously targets LECs and some non-lymphatic CNS compartments. With the surge of whole-organ and whole-body single cell/nucleus RNA sequencing, we reasoned that it may be possible to identify a marker enriched in LECs that would spare another CNS compartments. Mining our single-cell transcriptomic atlas of the dura, we identified Mmrn1 as the most significantly enriched transcript in dural LECs. Mmrn1 encodes multimerin 1, a large soluble homomultimeric adhesive glycoprotein^34,35^ first characterized in platelets where it acts as a carrier for coagulation factor V in both human and mouse^36,37^. Expression has also been reported in human endothelial cells^38^ and was described as enriched in human lymphatic endothelial cells^39^. These properties made Mmrn1 an attractive candidate on which to build a lymphatic Cre driver.

Here we generated an Mmrn1-T2A-CreERT2 knock-in mouse and tested its recombination profile across the CNS and the periphery. In adult mice, Mmrn1CreERT2 drives recombination throughout the lymphatic network with no recombination in non-LECs in the CNS, or in the periphery. When induction is moved into the postnatal window during which dural lymphatics develop, specificity is largely maintained, with a small population of recombined brain cells, predominantly Glut1+ endothelium, representing less than 0.5% of the brain vasculature. The Mmrn1CreERT2 line therefore provides a lymphatic driver with adult CNS specificity that, to our knowledge, no existing tool offers.

## Results

### Mmrn1 is enriched in lymphatic endothelial cells in the CNS

Identifying cell-specific markers is paramount to generate tools to assess the role of specific cells in organ and system function. We therefore mined our published single cell-RNA sequencing of the mouse dural compartment^40^. LECs were identified among the dural cells by their expression of canonical markers Prox1, Flt4, Ccl21a and Lyve1 (Figure 1A, Supplementary Figure 1A and B). Generation of differentially expressed genes (DEGs) between LECs and other dural cells revealed that 25 genes were highly enriched in LECs (Figure 1B). Among these 25, canonical markers of LECs such as Prox1 and Ccl21 are among the most enriched, but Mmrn1 was found as the most significant and second most enriched gene in LECs (Figure 1B). UMAP representation of Mmrn1 expression in dural cells demonstrated confined expression within the LECs (Figure 1C). Mmrn1 is a gene encoding multimerin 1, a large adhesive glycoprotein^34^, with described function in blood coagulation^37^, and recently demonstrated to be enriched in human LECs^39^. First, we confirmed the transcriptomic enrichment of Mmrn1 in LECs. We analyzed the expression profile of Mmrn1 in the Tabula Muris single cell database^41^ (Figure 1D) and a recent organ-specific single-cell RNA sequencing dataset^42^ (Figure 1E). In both datasets, we found Mmrn1 enriched in LECs (Figure 1D and E), demonstrating that it may be a suitable marker for LECs. We then confirmed that, using a published RNA sequencing dataset^43^, and contrary to other known lymphatic markers, Mmrn1 appears to be virtually absent from brain cells under homeostatic conditions (Figure 1F),.

**Figure 1:**
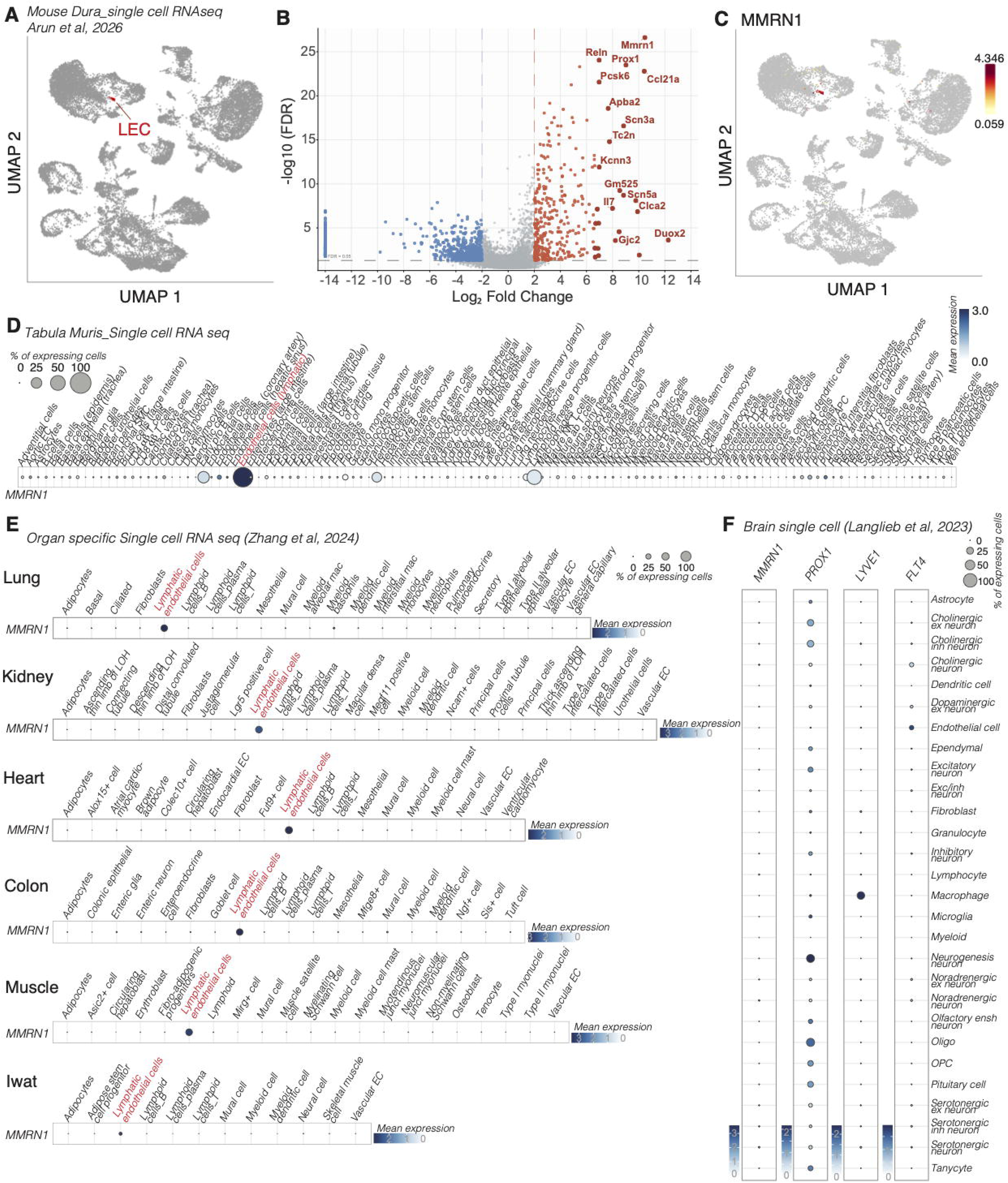
Mmrn1 transcript is enriched in lymphatic endothelial cells. A. UMAP plot of dural cells identifying the lymphatic endothelial cells (LECs, red) cluster from other dural cells based on canonical markers. B. Volcano plot of differentially expressed genes in dural lymphatic endothelial cells compared to other dural cells. Genes enriched in LECs are shown in red. C. UMAP plot of Mmrn1 expression in dural cells highlighting enrichment in LECs. D. Dot plot of expression of Mmrn1 in the Tabula Muris single-cell RNA sequencing dataset covering most organs of the adult mouse. The LECs cluster is highlighted in red. E. Dot plot of expression of Mmrn1 in single-cell RNA sequencing dataset of the lung, kidney, heart, colon, muscle and inguinal white adipose tissue (IWAT) of adult mouse. F. Dot plot of expression of Mmrn1, Prox1, Lyve1 and Flt4 in single-cell RNA sequencing dataset of the adult mouse brain.

Overall, these transcriptomic analyses suggest that Mmrn1 may be a valuable and specific marker for LECs with no detectable RNA expression in CNS cells.

### Generation and validation of expression of Mmrn1 in the CNS

To decipher if Mmrn1 could be a new marker for LECs in the CNS, we generated a Mmrn1CreERT2 mouse via insertion of the T2A-CreERT2 cassette between the last (8^th^) exon of Mmrn1 and its stop codon (Figure 2A). These mice were then crossed with the Ai6 stop flox^44^ mice to assess recombination specificity and efficacy. Mice at P30 were injected with tamoxifen at 45 mg/kg every other day for a total of two injections, and Ai6 reporter expression was analyzed at P60 (Figure 2B). Macroscopic imaging of the dorsal and ventral parts of the dura, still attached in their skull beds, demonstrated recombination in the Mmrn1CreERT2+ mice along the sinuses and the middle meningeal artery (arrows, Figure 2C). Recombination was absent in Mmrn1CreERT2-mice (Figure 2C), or in Mmrn1CreERT2+ mice not injected with tamoxifen (Supplementary Figure 1C). The fluorescence pattern traced a vascular structure consistent, both morphologically and anatomically, with dural lymphatic vessels. Whole mount preparation of the dorsal portion of the dura confirmed recombination in Lyve1+ lymphatic vessels (Figure 2D and E) with no recombination detected in Lyve1+ macrophages (Figure 2E; yellow arrow) or in any other structure outside of the lymphatic vessels (Figure 2D and E). We found similar lymphatic-specific recombination along the middle meningeal artery (Supplementary Figure 1D), at the base of the skull (Supplementary Figure 1E), and around the cribriform plate (Supplementary Figure 1F). The recombination in LECs was also demonstrated in the spinal cord meninges (Figure 2F and Supplementary Figure 1G). We measured the morphology of the dural lymphatics in CreERT2+ and CreERT2-mice, using the Lyve1 staining, to ensure that the Cre expression does not alter the lymphatics. We found no change in total lymphatic length, number of sprouts or lymphatic width, demonstrating that the presence of the Cre did not alter the morphology of the dural lymphatics (Figure 2G). We then analyzed the brain and spinal cord of these mice. Macroscopic imaging of the brain did not reveal any obvious recombined cells in the Mmrn1CreERT2+ mice compared to the Mmrn1CreERT2-(Figure 2H). Brain and spinal cord sections of Mmrn1CreERT2+ mice did not display any Ai6+ cells in either the brain or the spinal cord (Figure 2I and J), demonstrating that the Mmrn1-induced recombination is specific to the dural LECs without any observable recombination in other CNS cells.

**Figure 2:**
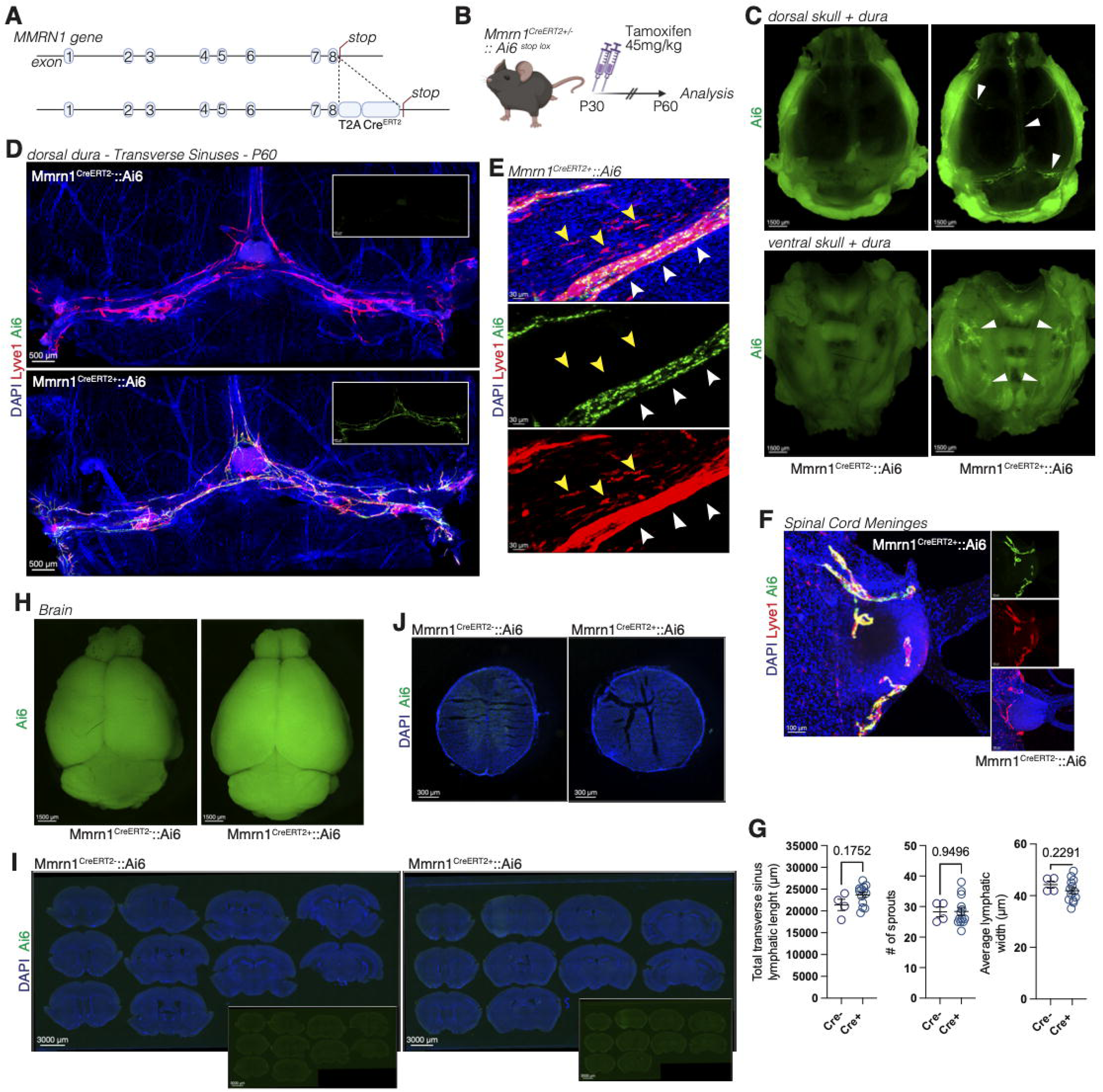
Mmrn1 CreERT2 recombination in adult mice is lymphatic-specific in the CNS. A. Scheme of the generation of the Mmrn1 CreERT2 mice. The T2A-CreERT2 cassette was inserted after the 8^th^ exon of the mouse Mmrn1 gene. Mice were then crossed with the Ai6 stop flox mice. B. Scheme of tamoxifen regimen. Mice at P30 were injected with tamoxifen for every other days (2 injections total) and analyzed a month later. C. Representative macroscopic images of the mouse dura, within the skull beds of Mmrn1CreERT2- and Mmrn1CreERT2+ mice. Arrows point to recombined lymphatic vessels. D. Representative confocal images of the dural lymphatic vessels (Lyve1, red) along the transverse sinus of Mmrn1CreERT2- and Mmrn1CreERT2+ mice. E. Representative images of the Ai6 recombination in the Lyve1+ lymphatic vessels (white arrow) but not Lyve1+ macrophages (yellow arrow) in the dura of Mmrn1CreERT2+ mice. F. Representative images of the Ai6 recombination in the Lyve1+ lymphatic vessels of the spinal cord meninges of the Mmrn1CreERT2+ mice. G. Quantification of the total lymphatic length, number of lymphatic sprouts, and average lymphatic width of the dural lymphatic vessels along the transverse sinuses of adult Mmrn1CreERT2- and Mmrn1CreERT2+ mice. Mean +/- s.e.m. Welch’s t test. H. Representative macroscopic images of the dorsal view of the whole brain of Mmrn1CreERT2-and Mmrn1CreERT2+ mice. I. Representative images of Ai6 recombination in the brain of Mmrn1CreERT2- and Mmrn1CreERT2+ mice. J. Representative images of Ai6 recombination in the spinal cord of Mmrn1CreERT2- and Mmrn1CreERT2+ mice.

### Mmrn1 CreERT2 mice demonstrate lymphatic-specific recombination in the periphery

To evaluate recombination specificity outside of the CNS, the recombination in several peripheral organs was assessed. Whole mounts of the ear and the diaphragm showed recombination in a pattern reminiscent of the lymphatic vasculature of these organs (Figure 3A and B). Confocal imaging of Ai6 recombination in the ear and diaphragm against the canonical lymphatic marker Lyve1 confirmed that the expression of Ai6 is confined within the Lyve1 expressing vessels with no detectable expression in non-lymphatics (Supplementary Figure 2A and B). The Ai6 expression appears to be independent of the level of Lyve1 expression by the lymphatic vessels (Supplementary Figure 2A and B). Macroscopic imaging of whole organs demonstrated recombination in the extensive lymphatic network of the heart (Figure 3C) and the lung (Figure 3D), with confocal confirmation of a restricted expression in Lyve1+ lymphatic vessels (Figure 3C and D). The gastrointestinal (GI) tract possesses an anatomically and functionally unique and complex network of lymphatic vessels^4,45^. We next examined the efficacy of our Cre driver to recombined in the GI tract. Macroscopic imaging of the mesentery highlighted Ai6 recombination of the collecting lymphatic vessels of the mesentery leading to the mesenteric lymph nodes (Figure 3E), a recombination that is absent in Cre-mice (Figure 3E). We can also identify strong Ai6 signal within the intestine itself (Figure 3E). We therefore sectioned the small intestine and found recombination of the Mmrn1CreERT2 in the both the lacteals (Figure 3F, white arrows) and in the lymphatic vessels of the submucosal plexus (Figure 3F, yellow arrow), with no recombination observed outside of the lymphatic vessels. These results demonstrate that the Mmrn1creERT2 mice allow broad and specific recombination in lymphatic beds across the entire mouse.

**Figure 3:**
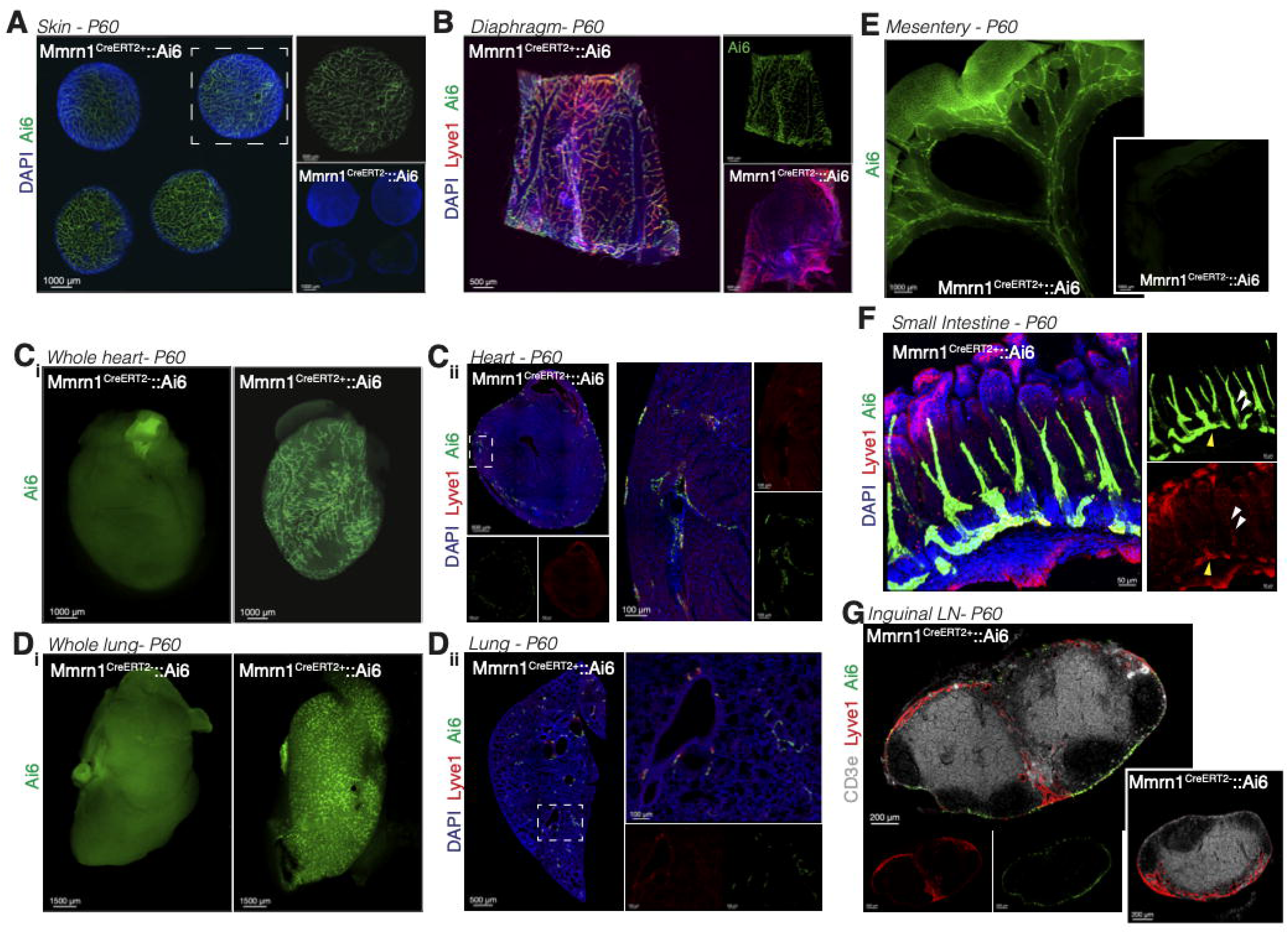
Mmrn1 CreERT2 recombination in peripheral organs. A. Representative images of the Ai6 recombination in the lymphatic vessels in the skin (ear) of Mmrn1CreERT2- and Mmrn1CreERT2+ mice. B. Representative images of the Ai6 recombination in the lymphatic vessels (Lyve1, red) in the diaphragm of Mmrn1CreERT2- and Mmrn1CreERT2+ mice. C. Representative macroscopic (i) and confocal (ii) images of Ai6 recombination in the lymphatic vessels (Lyve1, red) of the heart. D. Representative macroscopic (i) and confocal (ii) images of Ai6 recombination in the lymphatic vessels (Lyve1, red) of the lung. E. Representative macroscopic images of the Ai6 recombination in the lymphatic vessels of the mesentery of Mmrn1CreERT2- and Mmrn1CreERT2+ mice. F. Representative confocal images of the Ai6 recombination in the lacteals (Lyve1, red) in the small intestine of Mmrn1CreERT2+ mice. G. Representative confocal images of the Ai6 recombination in the LECs of the inguinal lymph nodes of Mmrn1CreERT2- and Mmrn1CreERT2+ mice.

Furthermore, lymph nodes have unique populations of LECs and serve specialized structural and immunological functions^46–49^. Analysis of recombination of Mmrn1CreERT2 mice in the inguinal lymph nodes showed Ai6 expression in the LECs of the capsule (Figure 3G, Supplementary Figure 2D), with only partial overlap with Lyve1 staining of the lymph node LECs. Independent analysis of a published single-cell RNA seq of the lymph node LECs^48^ identified 5 individual populations of LECs. Comparison of gene expression of Mmrn1 and Lyve1 in the different clusters of LECs demonstrated higher expression of Mmrn1 in the capsular LECs with low expression of Lyve-1. On the other hand, little to no expression of Mmrn1 was found in floor, MARCO+ (Medullary Sinus LECs), and transitional LECs, that express high levels of Lyve-1. Additionally, shared expression of Mmrn1 and Lyve1 was observed in Ptx3+ LECs (Central Medulla and Paracortical LECs) (Supplementary Figure 2C). Indeed, differentially expressed genes analysis characterizing the different LECs subtypes identified Mmrn1 as a marker for capsular lymph nodes LECs^48^. These results are concordant with our histological analysis that Mmrn1 is expressed highly in the capsular LECs with only partial overlap with Lyve1+ LECs. Overall, our data indicate that the Mmrn1CreERT2 mice allow specific recombination in lymphatic endothelial cells through the different organs of the mouse with LECs-subtype specific recombination in the lymph nodes LECs population.

### Recombination of Mmrn1CreERT2 during development of the dural lymphatics

We then sought to address if our new Cre system was specific and appropriate to study the development of the dural lymphatic system. We induced recombination with two tamoxifen injections (45mg/kg, every other day) starting at P10 to target the developing dural system^11,12^ (Figure 4A), and euthanized at P60. We found that similarly to the adult recombination, developmental induction of the Mmrn1CreERT2 leads to specific recombination in the lymphatic vasculature both using macroscopic imaging (Figure 4B) and confocal imaging (Figure 4C and D). Developmental recombination similarly did not induce major changes in dural lymphatic morphology (Supplementary Figure 3A), demonstrating the specificity and lack of toxicity of our system. We then analyzed both the brain and spinal cord for potential additional recombination in non-lymphatic structures.

**Figure 4:**
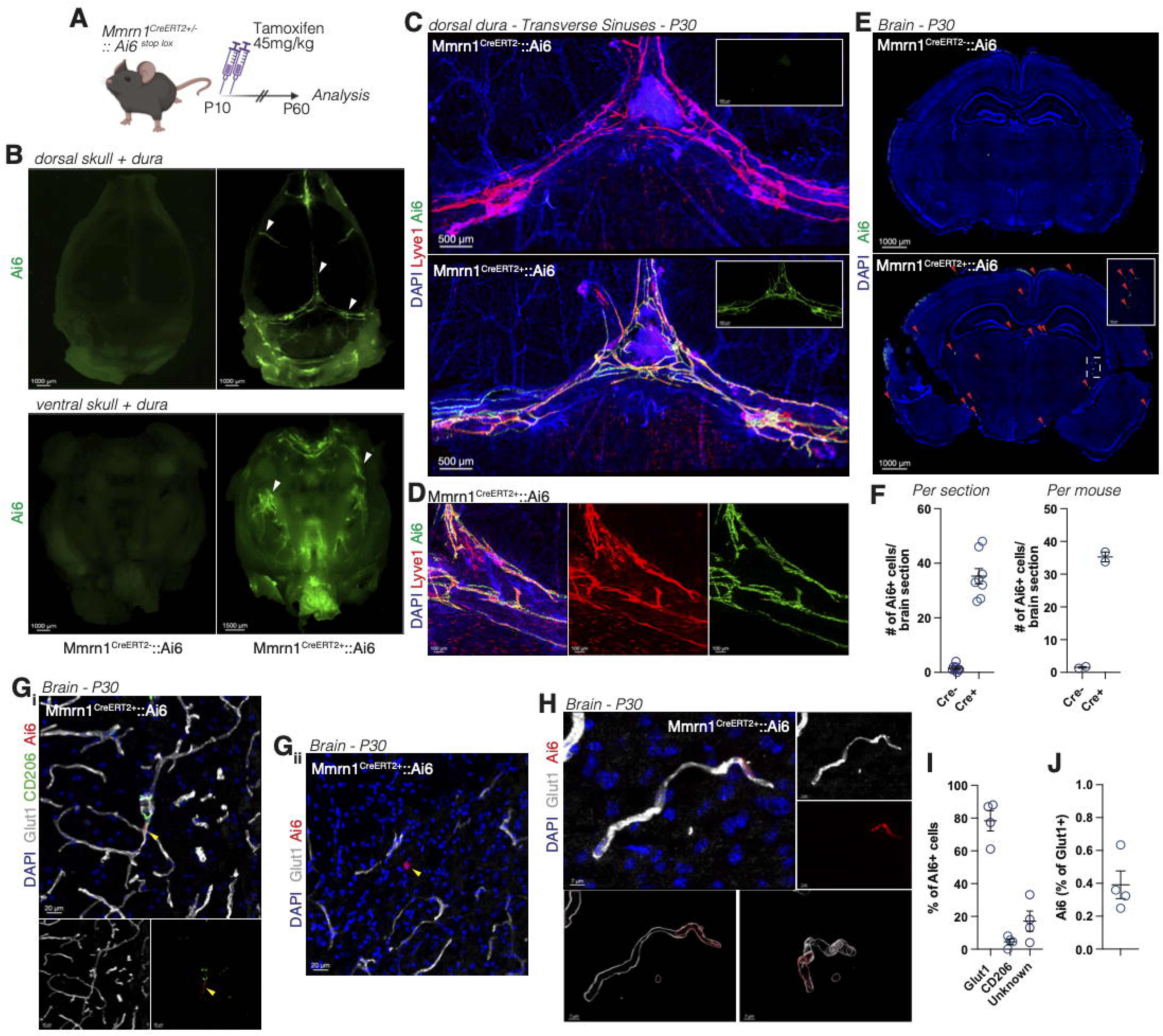
Analysis of Mmrn1CreERT2 recombination during dural lymphatic development. A. Scheme of tamoxifen regimen. Mice at P10 were injected with tamoxifen twice (every other day) and analyzed at P60. B. Representative macroscopic images of the mouse dura, inside of the skull beds of Mmrn1CreERT2- and Mmrn1CreERT2+ mice. Arrows point to recombined lymphatic vessels. C. Representative confocal images of the dural lymphatic vessels (Lyve1, red) along the transverse sinus of Mmrn1CreERT2- and Mmrn1CreERT2+ mice. D. Representative images of the Ai6 recombination in the Lyve1+ lymphatic vessels in the dura of Mmrn1CreERT2+ mice. E. Representative images of Ai6 recombination in the brain of Mmrn1CreERT2- and Mmrn1CreERT2+ mice. Red arrows point to Ai6+ cells. F. Quantification of the number of Ai6+ cells per brain section presented as average per brain section analyzed (4 individual sections per mouse) or pooled per mouse. Mean +/- s.e.m. G. Representative images of the Ai6 recombination in Glut1+ (i) or unknown (ii) cells in the brain of Mmrn1CreERT2+ mice. Yellow arrows point to Ai6+ cells. H. Representative 3D reconstruction of Ai6 recombination in Glut1+ endothelial cells in the brain of Mmrn1CreERT2+ mice. I. Quantification of the percentage of Ai6+ cells that colocalize with Glut1, CD206 or no marker in the brain of Mmrn1CreERT2+ mice. Analysis of 2 independent brain sections from 2 independent mice. Mean +/- s.e.m. J. Quantification of the percentage of Glut1+ area that is Ai6+ in the brain of Mmrn1CreERT2+ mice. Analysis of 2 independent brain sections from 2 independent mice. Mean +/- s.e.m.

Contrary to the adult recombination, we found sparse recombination in cells of both the brain and the spinal cord (Figure 4E, Supplementary Figure 3B). Quantification of the number of Ai6+ cells showed around 35 cells per brain section (Figure 4F). We then decided to identify what cells were recombined in the brain of our Mmrn1CreERT2 mice when induced during early postnatal development. We found that the majority of Ai6+ cells in the brain colocalized with the Glut1 marker of the vasculature (Figure 4G and H), with few cells colocalizing with the perivascular macrophage marker CD206 or no marker at all (Figure 4G and H). Quantification demonstrated that around 80% of the Ai6+ cells in the brain are Glut1+, less than 5% are CD206+ and around 15% are of unknown identity (Figure 4I). We found that the majority of the unknown Ai6+ cells were primarily located on the surface of the brain, likely in the pial layer of the meninges (Supplementary Figure 3C).

Given that the vast majority of Ai6+ cells were Glut1+, suggesting a blood endothelial cells identity^50^, we decided to quantify how much of the Glut1+ vasculature was recombined in our mice. We found that less than 0.5% of the Glut1 positive vasculature is Ai6+ in the Mmrn1CreERT2+ mice when induced at P10 (Figure 4J). This suggests that while Mmrn1CreERT2+ induces recombination in endothelial cells, the percentage of endothelial cell recombined is minor among the brain endothelial cells. Overall, these results suggest that, during the developmental window of the dural lymphatic system, the Mmrn1CreERT2 drives recombination in LECs but also in other cells, albeit at low level.

We similarly assessed the peripheral recombination after tamoxifen injection at P10. As with the adult recombination, we found lymphatic-specific recombination in the diaphragm (Supplementary Figure 3D), skin (Supplementary Figure 3E), the heart (Supplementary Figure 3F) and lung (Supplementary Figure 3G). The pattern of Ai6 expression in all peripheral organs analyzed appeared to be specific and consistent with a lymphatic pattern^11,51–54^ and was similar to the patterning observed after adult recombination. Analysis of Ai6 recombination in the lymph nodes showed, similarly to the adult recombination, expression in the capsule LECs with only partial overlap with the Lyve1+ LECs (Supplementary Figure 3H). Overall, our data demonstrate that Mmrn1CreERT2 at P10 is broadly active in all lymphatic beds analyzed with minor recombination in the brain and spinal cord vasculature.

## Discussion

Most causal cell-specific manipulations performed on meningeal lymphatic vessels to date have carried an unresolved ambiguity. The Cre drivers available to the field are built on Prox1, Lyve1 and Vegfr3, each of which is expressed, within the CNS and its borders, by cells that are not LECs^23–25,27,30,32^. Phenotypic analyses have had to either rely on genes that are not expressed by the non-LEC cells or concomitant use of multiple Cre drivers. Here we show that Mmrn1, the most significantly enriched transcript in the dural LECs, can be used to build a driver that resolves this ambiguity. In adult Mmrn1CreERT2 mice, recombination covers the entire dural lymphatic networks and the lymphatic beds of every peripheral organ examined, while remaining undetectable in other CNS cells. When induction is moved into the early postnatal developmental window, specificity is largely but not entirely preserved.

While Mmrn1CreERT2 mice present an LEC-specific recombination in the CNS of adult mice, we detect a low and sparse recombination primarily in brain and spinal cord vascular cells with developmental recombination. This highlights a potential time- and space-restricted expression of Mmrn1 in brain endothelial cells. One study has reported the localization of Mmrn1 in Weibel-Palade bodies of human endothelial cells in vitro^38^. It is therefore possible that Mmrn1 may be expressed by some endothelial cells during development. It is however interesting that the blood endothelium recombination would be restricted to CNS organs with no obvious similar phenotype observed in the periphery, suggesting that there may be something unique to the CNS blood endothelium during early postnatal development. Around 15% of the recombined cells in the developmental recombination remain unidentified and appear to be mostly restricted to the pial surface. Further investigation into the identity of these cells during development is necessary.

Our study identifies Mmrn1 as a new driver of genetic recombination in LECs but does not address the function of Mmrn1 in LECs. Work in platelets demonstrates a role of platelet-derived Mmrn1 in coagulation by facilitating platelet adhesion and storage of Factor V^37^. Given the importance of the extracellular matrix in the regulation of lymphatic function^55–58^, it is possible that Mmrn1 regulates fluid and immune cell drainage. Indeed, Emilin1, a protein of the same family as Mmrn1, has been shown to regulate valve formation, lymphatic structure and drainage^59,60^.

One limitation of the current study is that it does not test the specificity of recombination in disease conditions, particularly inflammatory conditions where Mmrn1 expression could potentially be induced in other cells than LECs. The Mmrn1CreERT2 line would therefore need to be re-validated in any disease model.

Overall, the Mmrn1CreERT2 line provides a new valuable tool that the field can use to assess adult CNS lymphatic loss- or gain-of function.

## Methods

### Mice

All mice used were on a C57Bl/6 background, bred in-house, and maintained under specific pathogen-free conditions on a 14/10h light/dark cycle with ad libitum access to food and water. The Mmrn1_T2A_CreERT2 mice were generated by Taconic and bred as heterozygous mice with the Cre carried on either male or female mice. All experiment presented here use mice heterozygous for the Cre. The Ai6 mice (B6.Cg-*Gt(ROSA)2CSor*^tm6(CAG-ZsGreen1)Hze^/J) were purchased from Jackson Laboratory (Cat # 007906). All experiments were approved by the Institutional Animal Care and Use Committee of the Cleveland Clinic Research.

### Tamoxifen treatment

Tamoxifen (Sigma-Aldrich; Cat # T5648) was dissolved in corn oil (Sigma-Aldrich; Cat # C8267) at a concentration of 10mg/ml. It was injected intraperitoneally twice (every other day) at a dose of 45 mg/kg at the indicated times.

### Tissue collection

Mice were euthanized by intraperitoneal injection of Euthasol (100mg/kg) and perfused with 0.1 M of PBS (10ml/mouse). The skull (both top and bottom), brain, spinal cord, nasal cavity, heart, lungs, diaphragm, ear skin, gut and lymph nodes (inguinal lymph nodes and mesenteric lymph nodes) were harvested and fixed in 4% Neutral Buffered Formalin (NBF; Thermo Fisher Scientific; SF100-4) for 24 to 72 hours depending on the tissue. To isolate the brain and the meninges, skin was removed from the head, and the muscles were stripped from the skull. After sectioning of the mandibles, and lateral cutting of the skull starting from the foramen magnum, the brain was dislodged from the skull. Nasal cavities were decalcified for 72h with decalcifying solution (Sigma-Aldrich; Cat # D0818). Tissue to be sliced was transferred to PBS with 30% sucrose (Sigma-Aldrich; Cat # S8501) for 72h or until the organ sank. Tissues were then frozen in OCT (Thermo Fisher Scientific; Cat # 23-730-571) and cryosectioned at 40µm (Leica Microsystems) on glass slides (Fisherbrand Superfrost Plus; Cat # 12-550-15).

### Immunohistochemistry

Tissues were incubated with PBS containing 2% of normal serum (Goat: Abcam; Cat # ab7481; Chicken: Jackson ImmunoResearch; Cat # 003-000-120), 1% bovine serum albumin (BSA; Equitech Bio; Cat # BAH70), anti-CD16/32 (1/1000; BioXCell; Cat # BE0307), 0.1% Triton-X-100 (Fisher Scientific; Cat # BP151-100) and 0.05% Tween 20 (Fisher Scientific; BP337-100) for 1h at room temperature (RT). Subsequently, the tissues were incubated with appropriate dilutions of primary antibodies O/N at 4C in PBS containing 1% BSA and 0.5% Triton-X-100. Primary antibodies used are Rabbit anti-Lyve1 (Angiobio; Cat # 11-034, 1/500); Rat anti-Lyve1 eF660 (Thermo Fisher; Cat # 50-0443-80, 1/200); Rat anti-CD3e biotin (Thermo Fisher; Cat # 13-0032-82, 1/200); Rat anti-CD206 (BioRad; Cat # MCA225, 1/200); Rabbit anti-Glut1 (Abcam; Cat # ab15309, 1/500) and Goa anti-Iba1 (Abcam; Cat # ab5076, 1/500). Tissues were then washed 3 times for 5 minutes at RT with PBS followed by incubation with fluorescently conjugated secondary antibodies, or conjugated streptavidin (Jackson ImmunoResearch; Thermo Fisher Scientific) at 1/500 for 2h at RT in PBS with 1% BSA and 0.5% Triton-X-100. After 5 min incubation with DAPI (1µg/ml; Thermo Fisher; D1306), the tissues were washed 3 times for 5 min with PBS. The dura was dissected from the top of the skull as previously described^8^. Tissues were then mounted with Aqua-Mount (Fisher Scientific; 14-390-5).

### Imaging and analysis

Macroscopic imaging of whole organs was performed using a Leica THUNDER Imager (Leica Microsystems) with a 1x objective. Confocal images were acquired with a Leica Stellaris Confocal System (Leica Microsystems) using LAS X software. Images were acquired with a 10x or 40x objective with a resolution of 1024 x 1024 pixels with a z-step of 2 to 4µm. Quantification of lymphatic morphology was performed as previously described^11^. Quantification of the number of Ai6+ cells in the brain of Mmrn1CreERT2+ mice was performed using the multi-point function of Fiji ImageJ software (NIH). The percentage of Ai6+ that are Glut1+, CD206+ or unknown was manually counted based on marker co-expression using Fiji ImageJ software. The coverage of Glut1+ area by Ai6 was measured using Fiji ImageJ software. The Glut1 ROI was created based on the threshold of the Glut1 staining. The area of AI6 within the ROI was then measured, and the coverage was calculated by dividing the area of Ai6 by the area of Glut1 staining multiplied by 100 for percentage coverage.

### Analysis of RNA sequencing

All analyses were performed using BioTuring BBrowser Pro (BioTuring Inc., San Diego, CA, USA). For our own dataset, normalized and scaled gene expression data were used for principal component analysis (PCA), and the top principal components were used to construct a shared nearest neighbor (SNN) graph for unsupervised clustering. Cell clusters were visualized using Uniform Manifold Approximation and Projection (UMAP). Cell type identities were manually assigned to each cluster based on the expression of established marker genes for known dural cell population. For LEC-specific genes, DEGs were called using Venice statistical test and filtered based on log2 foldchange (above 2) and false discovery rate (below 0.05). For publicly available datasets, they were imported from the Talk2Data database into the BioTuring BBrowser Pro platform for gene expression analysis. The authors’ cell identification was used.

### Statistics

Sample sizes were chosen based on standard power calculations using similar published experiments. Statistical test usage is described in each figure, with the numerical value of the statistical test displayed on the graphs. Statistical analyses were performed using Prism 10 (GraphPad Software Inc).

## Lead Contac

Requests for further information and resources should be directed to and will be fulfilled by the lead contact Antoine Louveau

## Resource Availability

This study generated the Mmrn1CreERT2 mice that will be shared via MTA upon request to the lead contact Antoine Louveau.

## Data and code availability

This study did not generate new data or code.

## Acknowledgments

We thank all the members of the Louveau lab for their valuable comments during the generation of this manuscript.

## Contributions

Concept: AL; Investigation: CG, GAT; Formal analysis: CG, GAT, JL, AL; Writing: CG, AL.

