## Supplementary Figures for "A new Mmrn1-CreERT2 Mouse Model for High-Fidelity Labeling of Adult Lymphatic Endothelium"

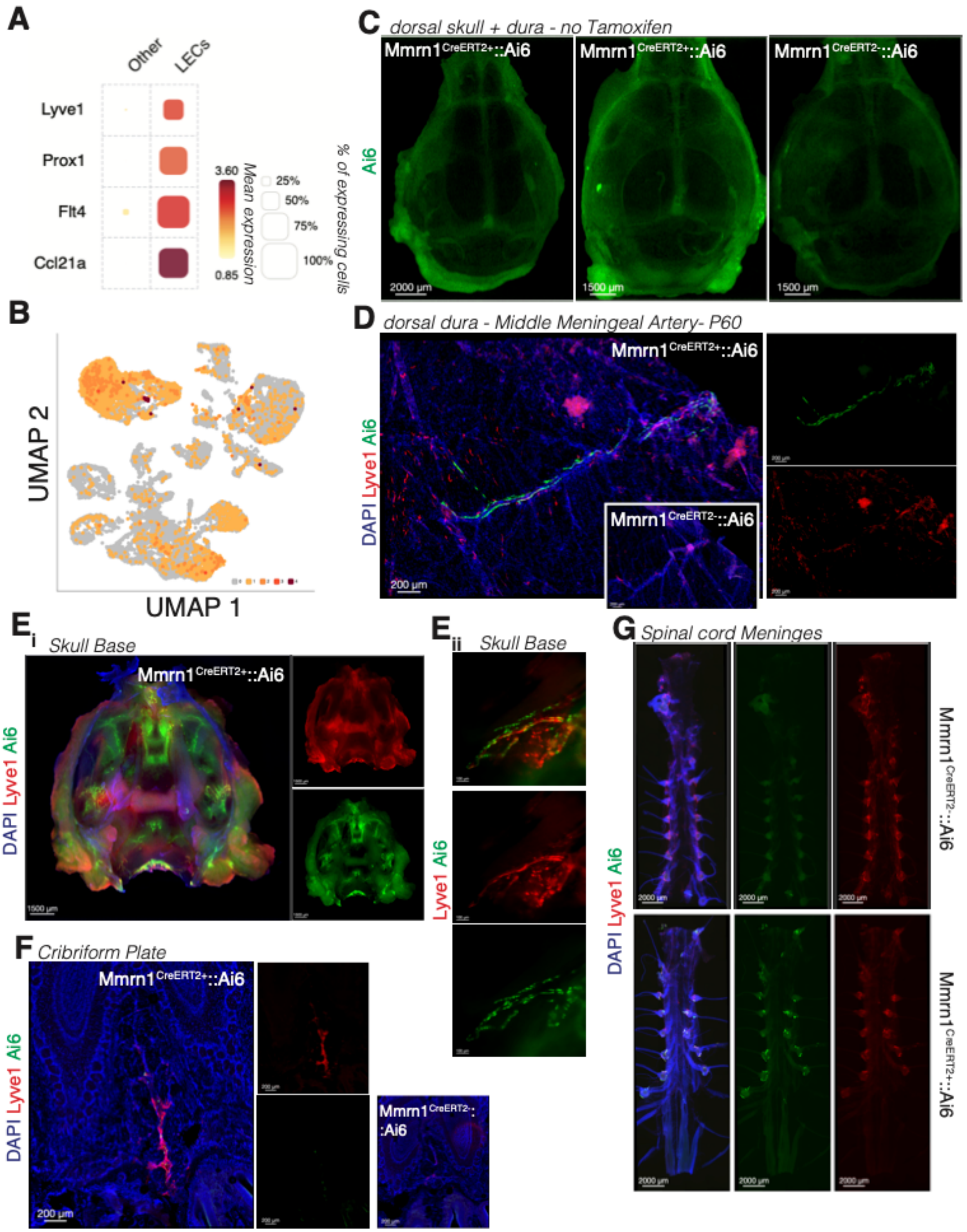

A
Other
LECs
Lyve1
Prox1
Flt4
Ccl21a
3.60
0.85
Mean expression
25%
50%
75%
100%
% of expressing cells
B
UMAP 2
UMAP 1
C
dorsal skull + dura - no Tamoxifen
Ai6
Mmrn1 CreERT2+::Ai6
Mmrn1 CreERT2+::Ai6
Mmrn1 CreERT2-::Ai6
2000 µm
1500 µm
1500 µm
D
dorsal dura - Middle Meningeal Artery- P60
DAPI Lyve1 Ai6
Mmrn1 CreERT2+::Ai6
Mmrn1 CreERT2-::Ai6
200 µm
Ei
Skull Base
DAPI Lyve1 Ai6
Mmrn1 CreERT2+::Ai6
1500 µm
Eii
Skull Base
Lyve1 Ai6
G
Spinal cord Meninges
DAPI Lyve1 Ai6
Mmrn1 CreERT2-::Ai6
Mmrn1 CreERT2+::Ai6
2000 µm
F
Cribriform Plate
DAPI Lyve1 Ai6
Mmrn1 CreERT2+::Ai6
Mmrn1 CreERT2-::Ai6
200 µm

**Supplementary Figure 1 (associated with Figure 1 and 2): Canonical LEC marker expression and Ai6 recombination in other CNS-lymphatic beds.**

A. Bubble plot of expression of canonical LECs markers in LECs cluster identified in Figure 1A compared to other dural cells.

B. UMAP of concomitant expression of canonical markers Lyve1, Prox1, Flt4 and Ccl21a in dural LECs.

C. Representative images of lack of Ai6 recombination in both Mmrn1CreERT2+ and Mmrn1CreERT2- mice when no tamoxifen is injected.

D. Representative images of Ai6 recombination in the lymphatic vessels along the middle meningeal artery in the dorsal portion of the dura of Mmrn1CreERT2- and Mmrn1CreERT2+ mice.

E. Representative images of Ai6 recombination in the lymphatic vessels (Lyve1, red) in the basal portion of the dura of Mmrn1CreERT2+ mice.

F. Representative images of Ai6 recombination in the lymphatic vessels (Lyve1, red) around the cribriform plate of Mmrn1CreERT2+ mice.

G. Representative images of Ai6 recombination in the lymphatic vessels (Lyve1, red) in the spinal cord meninges of Mmrn1CreERT2- and Mmrn1CreERT2+ mice.

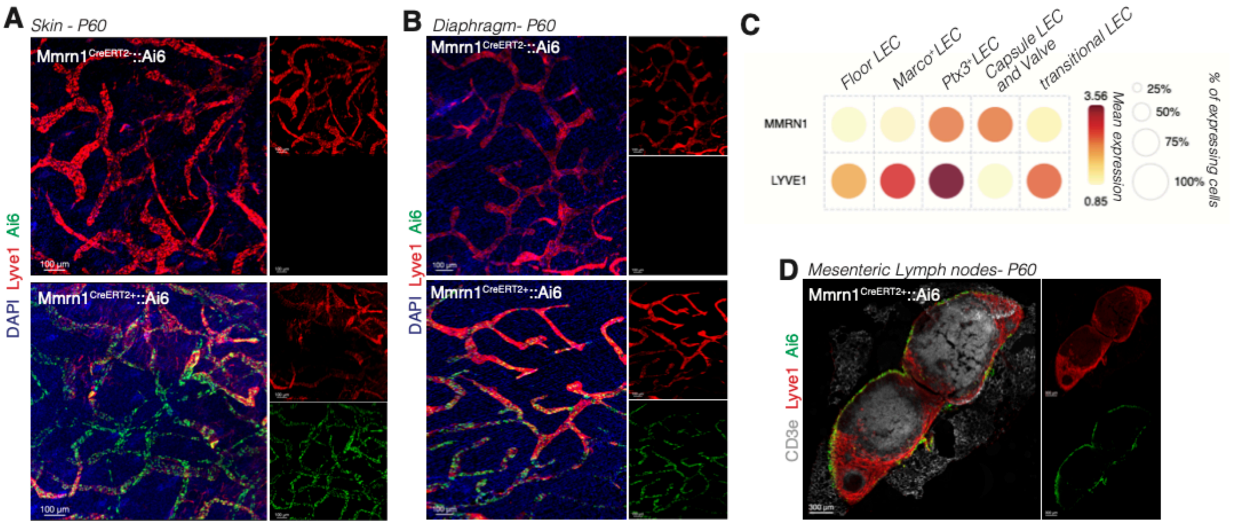
A
Skin - P60
Mmrn1 CreERT2-::Ai6
Mmrn1 CreERT2+::Ai6
DAPI Lyve1 Ai6
100 µm
B
Diaphragm- P60
Mmrn1 CreERT2-::Ai6
Mmrn1 CreERT2+::Ai6
DAPI Lyve1 Ai6
C
Floor LEC
Marco+LEC
Ptx3+LEC
Capsule LEC and Valve
transitional LEC
MMRN1
LYVE1
3.56
0.85
Mean expression
25%
50%
75%
100%
% of expressing cells
D
Mesenteric Lymph nodes- P60
Mmrn1 CreERT2+::Ai6
CD3e Lyve1 Ai6
300 µm

**<u>Supplementary Figure 2 (associated with Figure 3): Peripheral recombination of Mmrn1CreERT2+ mice.</u>**

A. Representative images of Ai6 recombination in the lymphatic vessels (Lyve1, red) in the skin (ear) of Mmrn1CreERT2- and Mmrn1CreERT2+ mice.

B. Representative images of Ai6 recombination in the lymphatic vessels (Lyve1, red) in the diaphragm of Mmrn1CreERT2- and Mmrn1CreERT2+ mice.

C. Bubble plot of transcript expression of Lyve1 and Mmrn1 in the lymph node LECs subsets.

D. Representative images of Ai6 recombination in the mesenteric lymph nodes of the Mmrn1CreERT2+ mice.

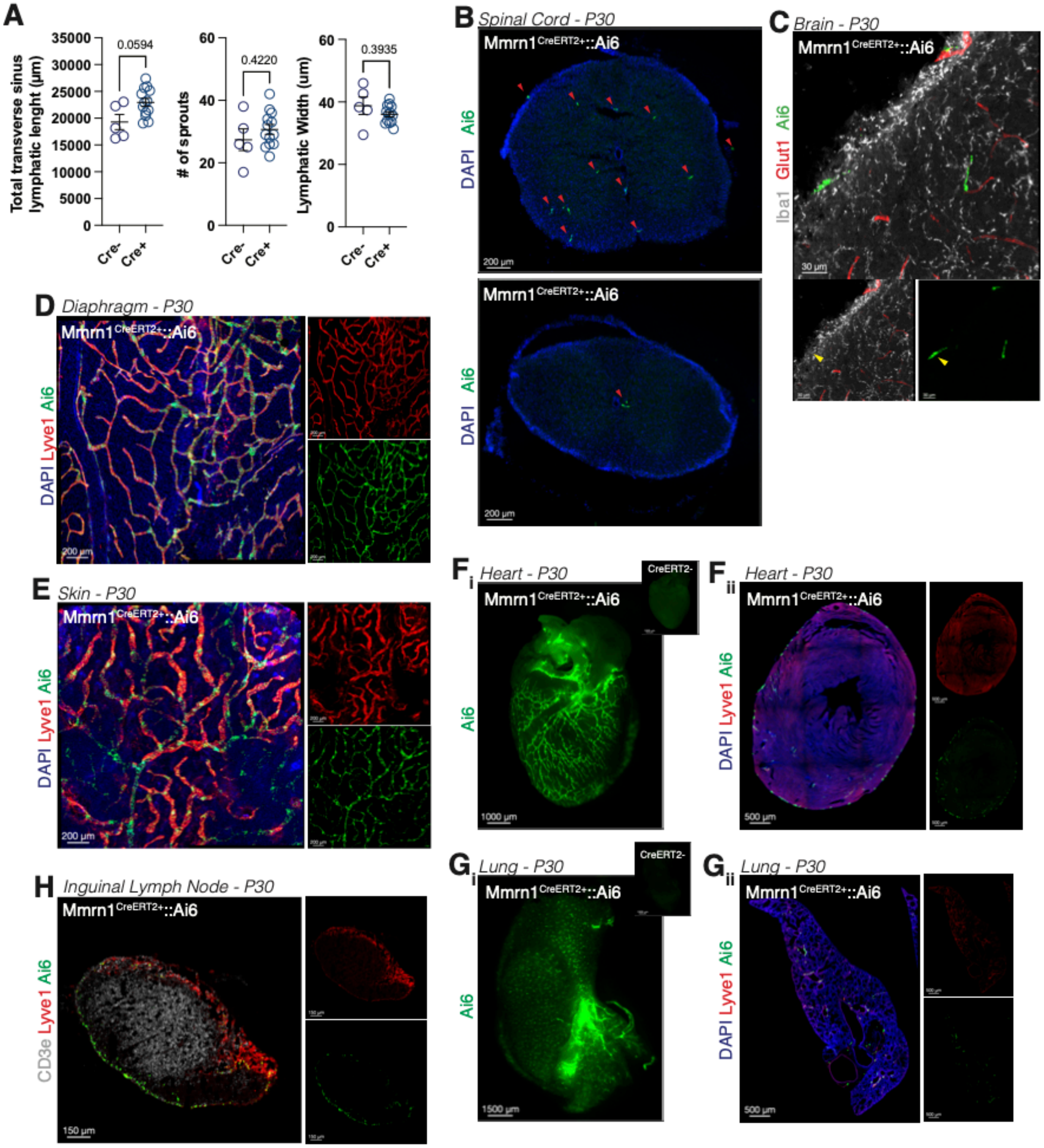


**Supplementary Figure 3 (associated with Figure 4): Developmental recombination of Mmrn1CreERT2 mice.**

A. Quantification of the total lymphatic length, number of lymphatic sprouts and average lymphatic width along the transverse sinus of Mmrn1CreERT2- and Mmrn1CreERT2+ mice after recombination at P10. Mean +/- s.e.m. Welch's t test.

B. Representative images of Ai6+ cells (red arrows) in the spinal cord of Mmrn1CreERT2+ mice.

C. Representative images of Ai6+ cells in the pial surface of Mmrn1CreERT2+ mice.

D. Representative images of Ai6 recombination in the lymphatic vessels (Lyve1, red) of the diaphragm of Mmrn1CreERT2+ mice.

E. Representative images of Ai6 recombination in the lymphatic vessels (Lyve1, red) of the skin (ear) of Mmrn1CreERT2+ mice.

F. Representative macroscopic (i) and confocal (ii) images of Ai6 recombination in the lymphatic vessels (Lyve1, red) of the heart of Mmrn1CreERT2- and Mmrn1CreERT2+ mice.

G. Representative macroscopic (i) and confocal (ii) images of Ai6 recombination in the lymphatic vessels (Lyve1, red) of the lung of Mmrn1CreERT2- and Mmrn1CreERT2+ mice.

H. Representative images of Ai6 recombination in the inguinal lymph node of Mmrn1CreERT2+ mice.
